# Changes in Ocular Fixation Characteristics Over Time during Reading

**DOI:** 10.1101/2025.06.09.658738

**Authors:** Lee Friedman, Oleg V. Komogortsev

**Affiliations:** Department of Computer Science, Texas State University, San Marcos, Texas, USA

## Abstract

In this report, we evaluate eye-movements during reading. There is a huge literature on this topic, but our report is not focused on the typical questions raised in this literature and our task design is very atypical for an eye-movement/reading study. While most reading studies evaluate mental processes during reading, the only mental process we evaluate is fatigue. While most reading studies use stimuli presented in the middle of the screen, our poem sections are presented in 24 lines of text that span the top to the bottom of the display. While most multi-line reading studies use a very large interline spacing, our interline spacing is more typical of text reading in newspapers, magazines and books. We report on changes in the characteristics of fixations as a function of time-on-task (TOT). We determine the start and end of reading for each subject/session and divide this reading time into 10 equal length (number of samples) periods (referred to as epochs). We looked for changes in fixation characteristics across epochs. We emphasize our results for horizontal position because these changes were monotonic and interpretable. For horizontal position signals, we found that the mean intersample distances in fixations increases, the rate of changes of fixation position decreases and the total fixation width increases as a function of epoch. We interpret our results to mean that, as reading progresses, it becomes more difficult to hold the eye still. Early in reading, the total fixation width is lower, the mean intersample distance is lower and there are frequent adjustments (i.e., changes in direction) to keep the eye well focused on the target word. As time progresses, it becomes more and more difficult to hold the eye perfectly steady, so fixations become wider and mean intersample distance increases.

## Introduction

There is a large literature involving the study of eye-movements during reading, including two impressive reviews by Rayner ([1], [2]). The present study is quite different from most of these studies. Most reading research has focused on understanding the mental processes that occur during reading [3]. The only “mental process” we investigate is fatigue, specifically, changes in fixation characteristics as a function of time on task (TOT). We are not aware of any prior study of changes in fixation characteristics as a function of (TOT) during reading.

According to Hyönä and Kaakinen [3]:

> “Reading texts longer than single sentences has been the least researched area in the study of eye movements during reading.”

Simalarly, Wang et al [4] state:

> “Over the past five decades, numerous empirical studies have argued that eye movements reflect moment-to-moment cognitive processing during reading. The evidence supporting this view largely derives from studies examining eye movements that occur during the reading of single-line sentences rather than multi-line texts, and this is the case even though people spend most of their time reading multi-line texts. In fact, we are consistently presented with multi-line texts from a variety of sources ranging from online materials such as advertisements and expository texts to novels and magazine articles.”

Our study involves reading six 4 line stanzas from a nonsense poem (“The Hunting of Snark” by Lewis Carroll, https://www.poetryfoundation.org/poems/43909/the-hunting-of-the-snark)

Also, from the Hyönä and Kaakinen paper [3]:

> “To date, readers’ eye movement recordings have been very productively put to use for studying word recognition, the size and nature of the effective visual field, and syntactic parsing of sentence and clause structure among competent adult readers.”

And

> “In a typical reading study, participants are asked to read single sentences silently for comprehension. Fixation durations on words and saccade lengths are then computed to examine how different word characteristics, which can be experimentally varied, influence the eye movement patterns.”

The present study does not address any of these topics.

And also from Hyönä and Kaakinen [3], for multi-line reading studies:

> “…the line spacing is adjusted to be wide enough (i.e., 2.5 line spacing) in order to reliably differentiate which line of text the reader is currently fixating on.”

In our case, the line spacing was much narrower and more typical of published text passages. We don’t think that this difference would affect our analysis. Also from Hyönä and Kaakinen [3]:

> “After reading, a measure of comprehension (e.g. free recall) is collected to check that the reader was engaged with the reading task; …”

In our case, the nonsense aspect of our poem might frustrate an analysis of comprehension.

Our poem stanzas are presented as a full page of text, from the top of the monitor screen to the bottom, whereas most reading research is focused on presenting text in the middle of the screen. Therefore, in our case, reading progress is potentially confounded with vertical position. We attempt to address this issue in our analysis.

All of the unusual features of our reading task stem from the fact that it was not designed to evaluate the typical objectives of eye-movement reading research. However, with some adjustments to our analysis, we believe that we can make certain statements about changes in fixation characteristics during reading as a function TOT. We employ a task collected more than ten years prior to the submission of this article. That database consists of more than 300 subjects tested on multiple occasions, and it was based on recordings from a relatively high-quality recording device (EyeLink 1000).

## Materials and methods

### The Eye Tracking Database

The eye tracking database employed in this study is fully described in [5] and is labeled “GazeBase” It is publicly available (https://figshare.com/articles/dataset/GazeBase_Data_Repository/12912257). The data were accessed on April 12, 2024. The authors had no access to information that could identify individual participants during or after data collection.

All details regarding the overall design of the study, subject recruitment, tasks and stimuli descriptions, calibration efforts, and eye tracking equipment are presented there. There were 9 temporally distinct “rounds” over a period of 37 months, and round 1 had the largest sample. For the present study, we only analyzed round 1 (322 subjects). There were were 151 females and 171 males. Data collection for Round 1 started in September, 2013 and ended in February of 2014. In each round, subjects were tested twice, with sessions typically approximately 20 min apart. Subjects were initially recruited from the undergraduate student population at Texas State University through email and targeted in-class announcements. Each session consisted of seven tasks. The main task task employed in the present study was the text reading task. Subjects were instructed to silently read a passage from the poem “The Hunting of the Snark” by Lewis Carroll. The six stanzas presented in session 1 are in Appendix Fig. 1. The six stanzas presented in session 2 are in Appendix Fig. 2. The task was automatically terminated after 60 seconds irrespective of the subjects’ reading progress. Only data collected during the first reading of the poem was included in our analysis. This reading period, the length of which was subject and session dependent, was divided up into 10 equal length (number of samples) sections, labeled epochs.

We also employed the a random saccade task. During the random saccade task, subjects were to follow a white target on a dark screen as the target was displaced at random locations across the display monitor, ranging from ± 15 and ± 9 of degrees of visual angle (dva) in the horizontal and vertical directions, respectively. The minimum amplitude between adjacent target displacements was 2 dva. At each target location, the target was stationary for 1 sec. There were 100 target positions per task (100,000 samples per task). The target positions were randomized for each recording. The distribution of target locations was chosen to ensure uniform coverage across the display.

Monocular (left) eye movements were captured at a 1,000 Hz sampling rate using an EyeLink 1000 eye tracker (SR Research, Ottawa, Ontario, Canada). The gaze position signals for the both tasks were classified into fixations, saccades, post-saccadic oscillations (PSOs) and various forms of artifact, using an updated version of our previously reported eye-movement classification method [6].

### Determining the Start of Reading

Subjects presented with our poems typically did not immediately begin reading. To find the start of reading, we searched for the first set of 4 fixations that were progressively more to the right of each other. The first fixation of the set was taken as the start of reading. The median time to start reading was 1.57 seconds after the poem display (25th to 75th % = [0.95 to 2.23 sec]). Typically, subjects began reading in the 8th fixation (25th to 75th % = [6th to 11th fixation]).

### Determining the End of Reading

If one simultaneously displays the entire horizontal and vertical position for the entire 60 sec duration of our task, it was possible to subjectively determine the point of the end of reading. The first author made this determination for all subjects.

### Calculation of Fixation Characteristics

For each fixation, we calculate the following measures:

1. The epoch the fixation belongs to.
2. The Mean Intersample Distances (horizontal and vertical)
3. The Percent Change Direction (horizontal and vertical)
4. The Fixation Width (horizontal and vertical)

#### Calculation of Mean Intersample Distance

If X is the position signal (horizontal or vertical), then:

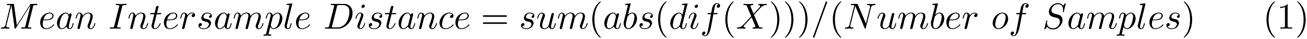

where *dif* is the difference operator. It calculates differences between adjacent elements of X. *abs* means absolute value. *sum* calculates the sum of the absolute value of the differences. A figure illustrating a fixation with low Mean Intersample Distance is presented in Fig. 1. A figure illustrating a fixation with high Mean Intersample Distance is presented in Fig. 2.

**Fig 1.**
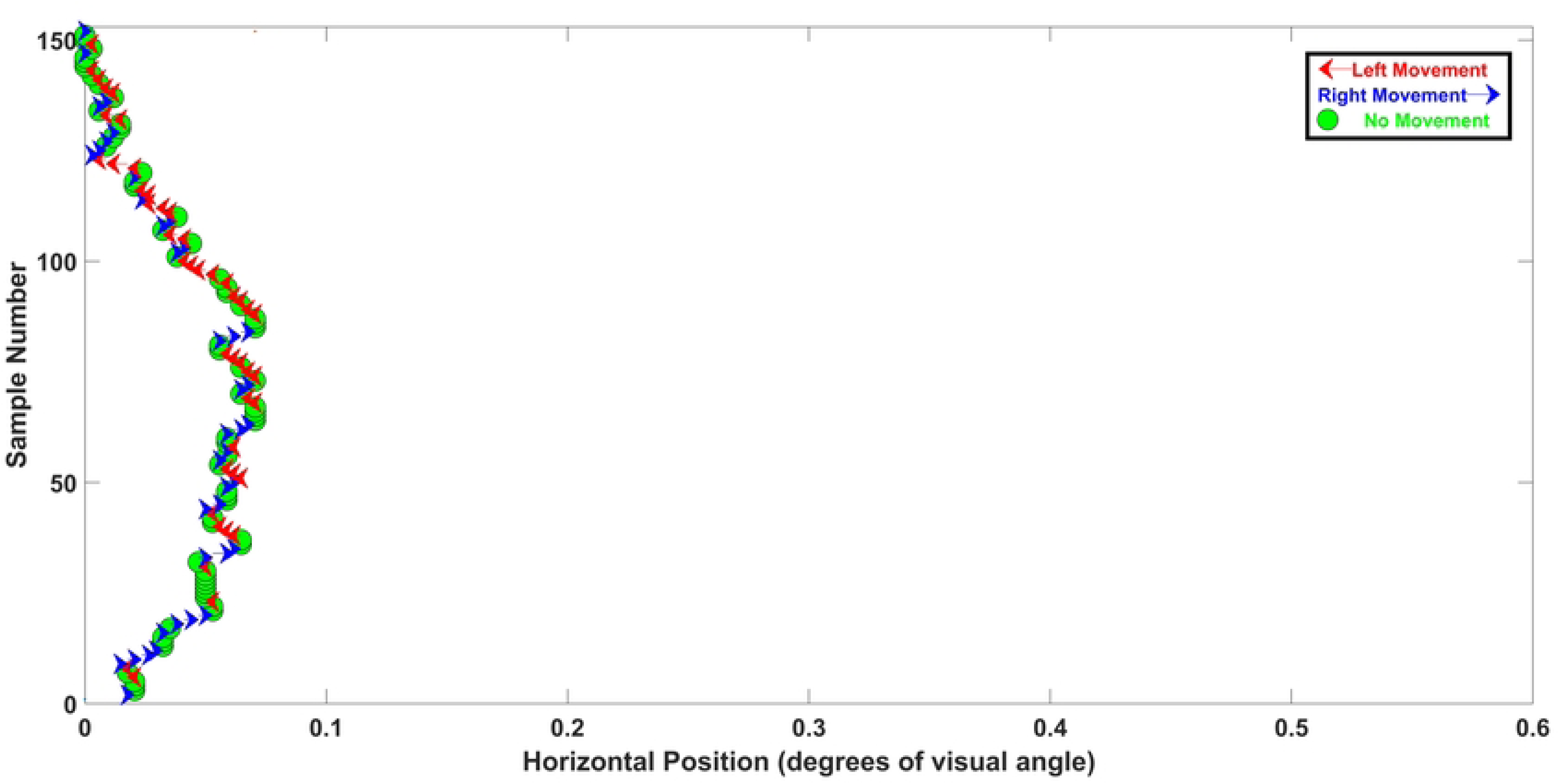
Illustration of the Percent Change Measure and the Mean Distance measure for an early fixation. The abscissa is horizontal position (shifted to start at 0.0, and set to a maximum of 0.6 dva to enhance the comparison with Fig. 2. The ordinate is Sample Number. This fixation is 153 ms long. The Percent Change value for this fixation was 55%, which is relatively high. The mean distance was 0.0021 dva, which is relatively low. As will become clear below, it is reasonable to think of this fixation as an archetype of a fixation early in the reading task, when fatigue has not set in. It has a very low mean intersample difference, a high percent change and a low fixation width.

**Fig 2.**
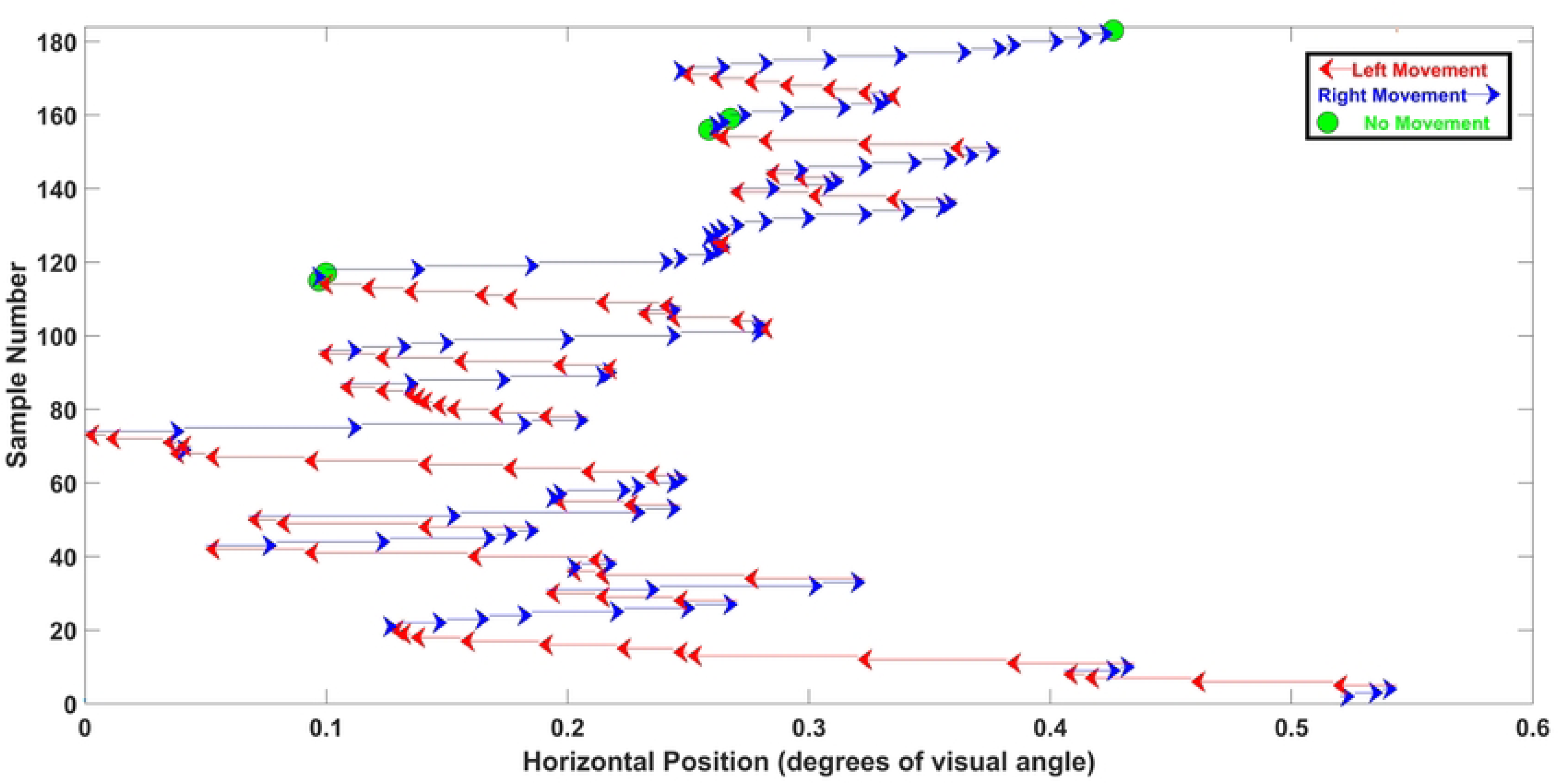
Illustration of the Percent Change Measure and the Mean Distance measure for a late fixation. The abscissa is horizontal position (shifted to start at 0.0). The ordinate is Sample Number. This fixation is 183 ms long. The Percent Change value for this fixation was 24.7%. The mean intersample distance was 0.0216 dva. As will become clear below, it is reasonable to think of this fixation as an archetype of a fixation late in the task, when fatigue has set in. It has a very high mean intersample difference, a low percent change and a high fixation width.

#### Calculate of Percent Change Direction

The first step in the calculation of Percent Change Direction is to create a vector of the signs (positive, no sign change or negative) of the differences between adjacent position samples (X) (“SignChange”):

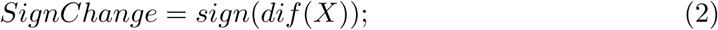

where dif is the difference operator. *sign* returns only the sign (+,0,-) of each difference. So, “SignChange” is a vector, (length number of samples −1) of sign changes over either the horizontal signal or the vertical signal. An example of a piece of a SignChange Vector is provided in Table 1

**Table 1.** Illustration of the Calculation of Percent Change Direction.

| SignChange | -1 | 1 | 1 | 0 | -1 | -1 | 1 | -1 | -1 | 0 | 1 | 1 | 1 | 1 | 1 |
| --- | --- | --- | --- | --- | --- | --- | --- | --- | --- | --- | --- | --- | --- | --- | --- |
| Count | 1 | 2 | 3 | 4 | 5 | 6 | 7 | 8 | 9 | 10 | 11 | 12 | 13 | 14 | 15 |
| Sum | 0 | 1 | 1 | 2 | 3 | 3 | 4 | 5 | 5 | 6 | 7 | 7 | 7 | 7 | 7 |

Next we sum the number of times that there is a direction change from the current SignChange value and the previous value. The 1st value is −1 and the 2nd value is 1, so at this point the sum is 1. The 3rd point is also 1, so the sum at this point is still 1. The 4th value is 0, which is counted as a sign change from 1, so at this point the sum is equal to 2. The 5th value is −1, and since this is a change from the prior value (0), the sum is now 3. The 6th value is also −1, so there is no change to the sum. However, the 7th value is 1, and since this is difference from the 6th value, the sum is now 4, etc… Over this 15 sample SignChange Vector, there are 7 direction changes, so the Percent Change Direction for the data in Table 1 is 46.67%. An example of a fixation with a high Percent Change Direction is presented in Fig. 1. An example of a fixation with a low Percent Change Direction is presented in Fig. 2.

### Calculation of Fixation Total Width

If our position signal (horizontal or vertical) for a fixation is X, then Fixation Total width is:

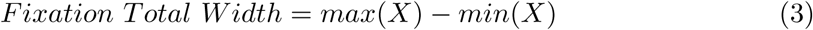

### Evaluating the Role of Vertical Position on Key Measures

As a subject proceeds to read our poem sections, the vertical position of the text moves from high (+10 dva) to low (−10 dva). Our focus is on effects of time on task but there is the logical possibility that the changes we see in fixation characteristics could simply reflect the vertical position of the visual stimulus. To check this, we used our random saccade task, described above. We divided all fixations from that task into 10 separate vertical position intervals (Table 2).

**Table 2.** Table of Divisions of Vertical Position in Degrees of Visual Angle.

| Interval | 1 | 2 | 3 | 4 | 5 | 6 | 7 | 8 | 9 | 10 |
| --- | --- | --- | --- | --- | --- | --- | --- | --- | --- | --- |
| Degrees | >8 | 6 ≤v<8 | 4 ≤v<6 | 2 ≤v<4 | 0 ≤v<2 | -2 ≤v<0 | -4 ≤v<-2 | -6 ≤v<-4 | -8≤v<-6 | v≤-8 |

We measured our all of our fixation characteristics in each of these vertical position bands. When we plot our results for reading, the x-axis is “Epochs”, which divide the reading time into 10 equal length periods, but we also plot our results for the 10 vertical position bands on the same plot, with epoch referring to vertical position. This allows us to evaluate the extent to which our results regarding time on task might simply reflect vertical position.

### Statistical Analyses

#### Fit of line to Medians

For the analysis of each measure, we first present a box-plot which includes the inter-quartile ranges and the medians across epochs. Next we fit an exponential proportional rate growth (or decrease) curve across the ten epochs (X=1 to 10). In all cases but 1, the formula for this curve was:

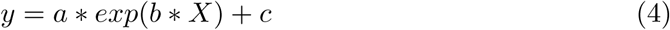

In one case (Mean Intersample Distance, (Vertical)), the form of the equation we fit was

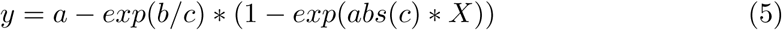

For each measure, we present the final fit of the equation with the estimates of *a, b, and c*. We also present the model *R*^2^.

#### Statistical Analysis using the Kruskal-Wallis test

The Kruskal-Wallis test is a non-parametric test and is analogous to the one-way analysis of variance (ANOVA). We tested the effect of Epoch. Each test provides a *χ̃*^2^ which is analogous to the *F* − *value* for a one-way ANOVA. Each test had a numerator degree of freedom (df) = 9 and a denominator df =116,656. Each test also provided a *p* − *value*. The *χ̃*^2^ and the *p* − *value* is presented in the relevant figure. These tests are based on the ranks of the observations.

Post-hoc tests, controlled for multiple comparisons using the Tukey Honestly Significant Difference procedure, were conducted. In most cases, the majority of epoch vs epoch comparisons were statistically significant. In such cases, we listed the non-statistically significant comparisons in the figure legend.

### Analysis of Word Length and Word Frequency as a Function of Word Number

We needed to know if the changes we see in our fixation characteristics were related to word length (number of letters) changes or word frequency (frequency of occurrence in the language) changes as a function of word number. The measure of word length (number of letters) is obvious. For our measure of word frequency we used the *SUBTLEX_US_*[7] database. We plotted log (base 10) Word Frequency against word number. We also fit at linear regression line to each such plot.

## Results

### Reading Time and Reading Speed

Our task was one minute in duration. Of 644 studies (322 subjects tests on two sessions), in only 449 studies did subjects finished reading the entire poem in 60 sec. All of our data presented below is based only on subjects who finished reading our poems in 60 sec or less. The median time to finish was 0.77 minutes (46.2 sec, Fig 3 (A)). Reading speed of subjects who finished reading the poems ranged from about 200 to nearly 550 words per minute (median 255.2 words per minute, Fig. 3 (B)).

**Fig 3.**
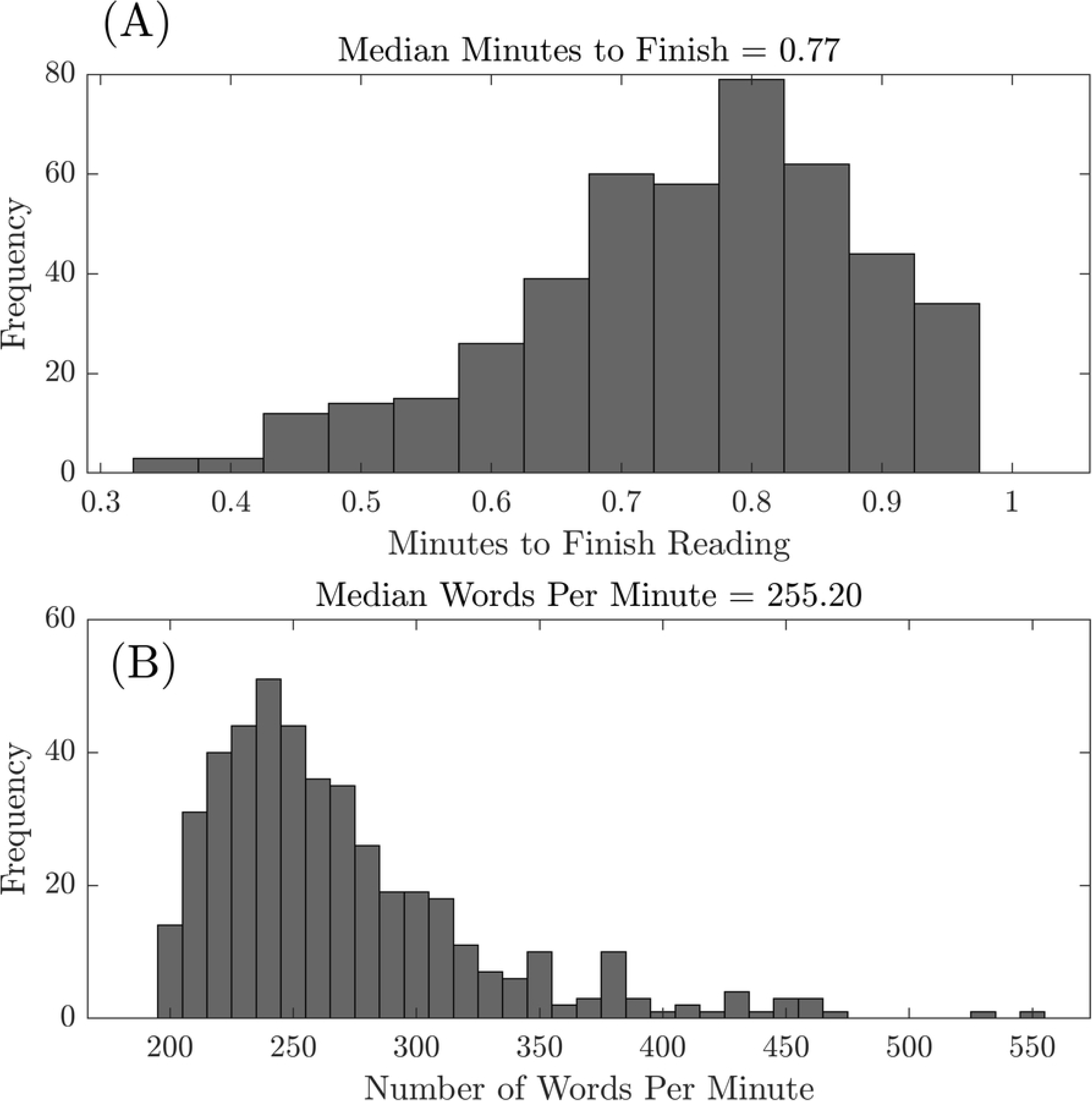
(A) Histogram of Reading Time. The task was 1 min in duration. Subjects who did not finish reading the poem in 1 min were excluded. This is the frequency distribution of minutes to finish for the remaining subjects. (B) This is the distribution of reading rate (words per minute) of the subjects who finished reading the poem.

### Number of Fixations Per Epoch and Number of Samples per Fixation per Epoch

Before detailing the changes we see in fixation characteristics with time-on-task in reading, we present some basic information about the number of fixations and fixation duration per epoch. In Fig. 4, we present a swarmchart of the number of fixations per epoch. Since these data are approximately normally distributed we describe the central tendency of the data per epoch with a mean rather than a median. Although there is evidence for a power-law-based decline across epochs, the range of means is narrow (17.4 to 19). The number of samples per fixation was also reasonably stable (Fig. 5). The range of median number of samples per fixation across epochs was 183 to 193 samples (median = 190.5 ms).

**Fig 4.**
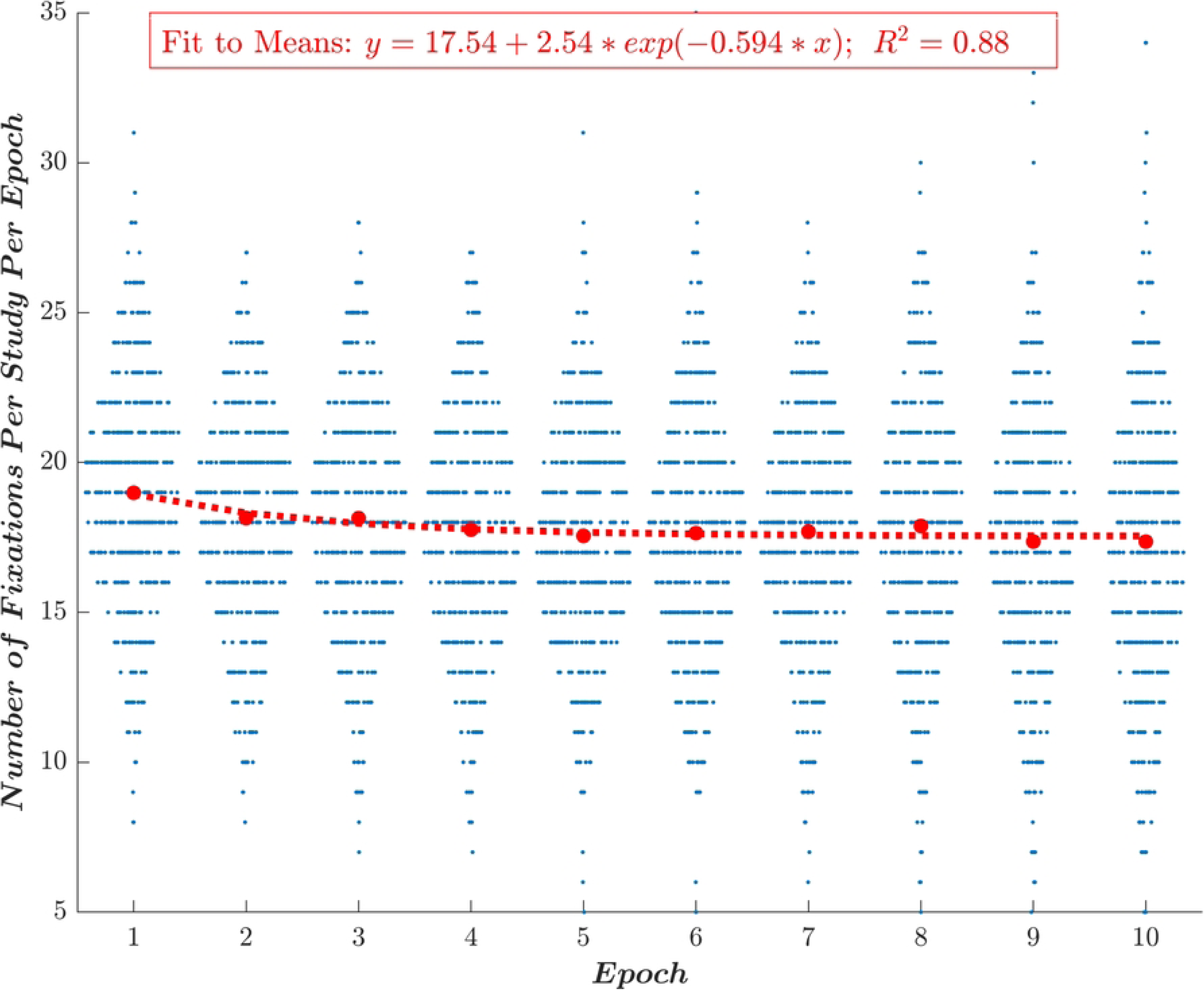
Swarmchart of the number of fixations per epoch. The red dots are the means. The red dashed line is the fit of expoential decay function to the means. The range of means is from 20.2 to 18 fixations per epoch. There is a gradual decline starting from the maximal value for epoch 1 to a relatively stable value for epochs 5 to 10.

**Fig 5.**
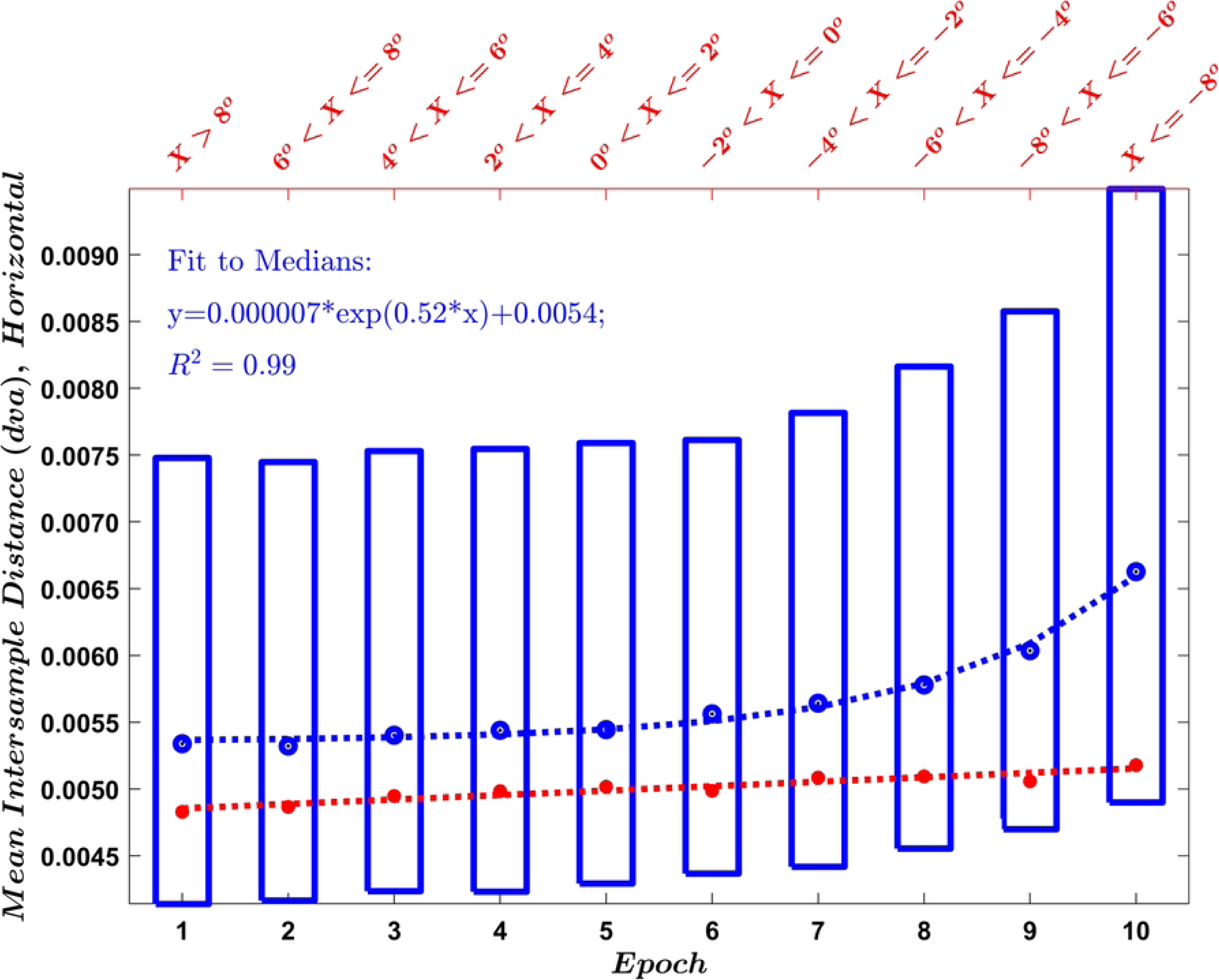
Boxplots of the number of samples per fixation across epochs. The blue boxes represent the results for all fixations. The bottoms of the blue boxes are the 25th percentiles. The tops of the blue boxes are the 75th percentiles. The blue dots are the medians. Samples are 1 ms apart.

### Horizontal versus Vertical Signal Results

We analyzed both the horizontal and vertical signals during each fixation. Although we found statistically significant changes in the vertical dimension, these results were not monotonic. Non-monotonic results require much more complicated explanations than monotonic results. Therefore, in the main report, we present only results for the horizontal dimension. We briefly present the vertical results in the Appendix (Appendix Figures 3, 4, 5).

### Mean Intersample Distance

The results for Mean Intersample Distance are presented in Fig. 6. The median Mean Intersample Distances (in blue) are relatively stable for epochs one to five. From then on, there is a gradual (exponential) increase in Mean Intersample Distance. The fit of the exponential proportional growth rate curve (blue dashed line) is good. The red line evaluates the effect of vertical position during the random saccade task. The red line is fit well to a regression line with a positive slope. The trajectory of the red line is very different from the trajectory of the blue line. We interpret this to mean that the results from reading (blue) are not simply a reflection of vertical position but rather are mostly an effect of time-on-task.

**Fig 6.**
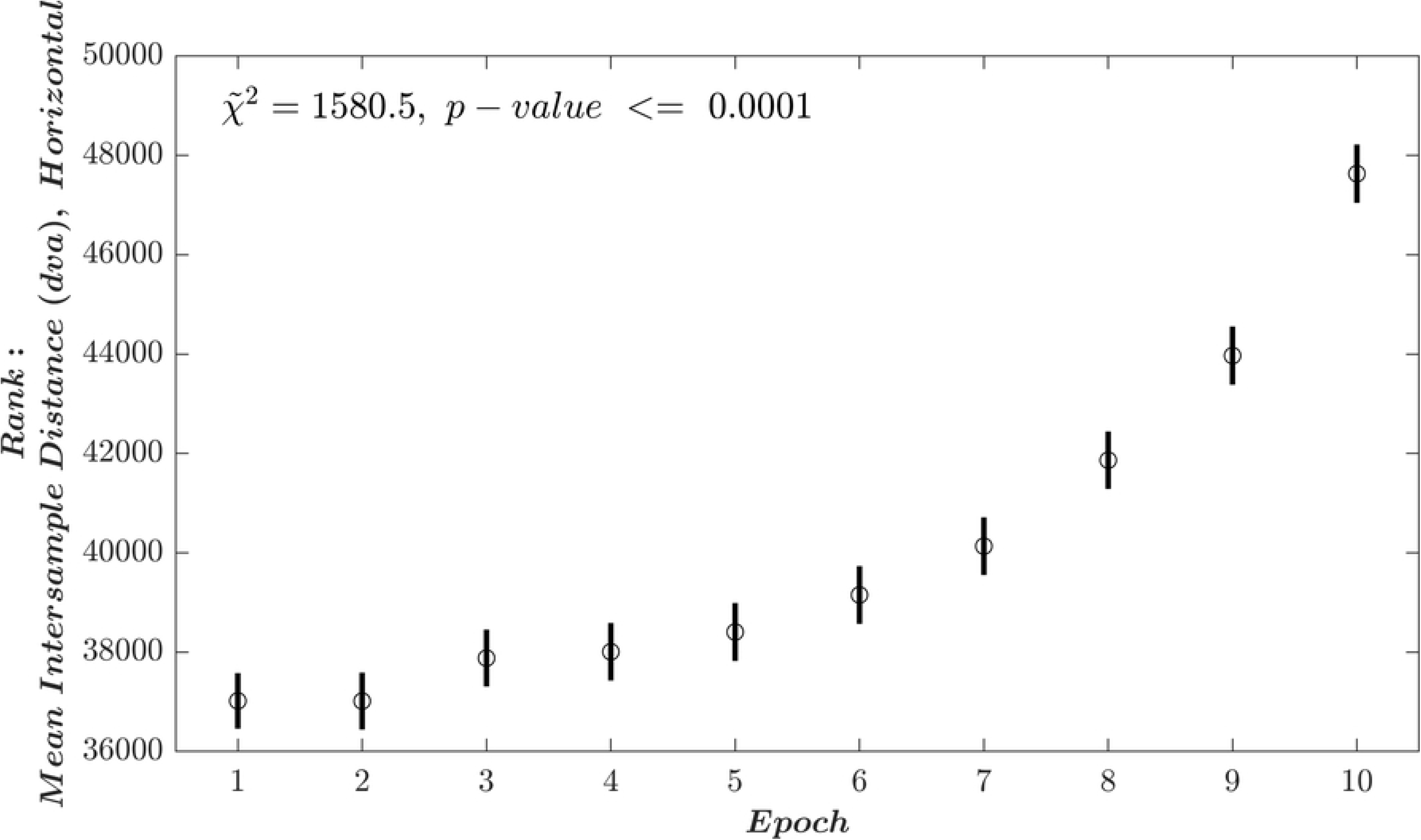
Results for Mean Intersample Distance (Horizontal). The results in blue emerge from analysis of the fixations during reading of the 6 poem stanzas (Appendix Figs. 1 and 2) from the TEX task. The relevant x-axis for the epoch (i.e., time) data is the bottom x-axis. The blue boxes represent the results for all fixations. The bottoms of the blue boxes are the 25th percentiles. The tops of the blue boxes are the 75th percentiles. The blue dots are the median Mean Intersample Distance for each epoch. The dashed blue line is the fit of a exponential proportional rate growth curve. In this case, the fit is excellent (*R*^2^ = 0.99). The red dots and red line are interpreted using the upper x-axis. These are from the random saccade task, not the reading task. The first red dot on the left represents the median Mean Intersample Distance when the mean eye position during fixations is >8°. The last red dot on the right represents the median Mean Intersample Distance when the mean eye position during fixations is <= −8°. The position data is fit well to a regression line with a positive slope (*R*^2^ = 0.90, *F* = 71.8, *p* − *value* < 0.0001, *df* = 1*/*9)

In Fig. 7, we present the statistical results comparing all the mean intersample distances for all fixations during each epoch. The *χ̃*^2^ is very large, meaning that there are definitely statistically significant differences in Mean Intersample Distance across epochs. The dots and bars are in units of rank and are the results of post-hoc comparisons between epochs. These comparison are corrected for multiple comparisons. If the bars for any two-epoch comparison do not overlap, this means that the epochs are statistically significantly different.

**Fig 7.**
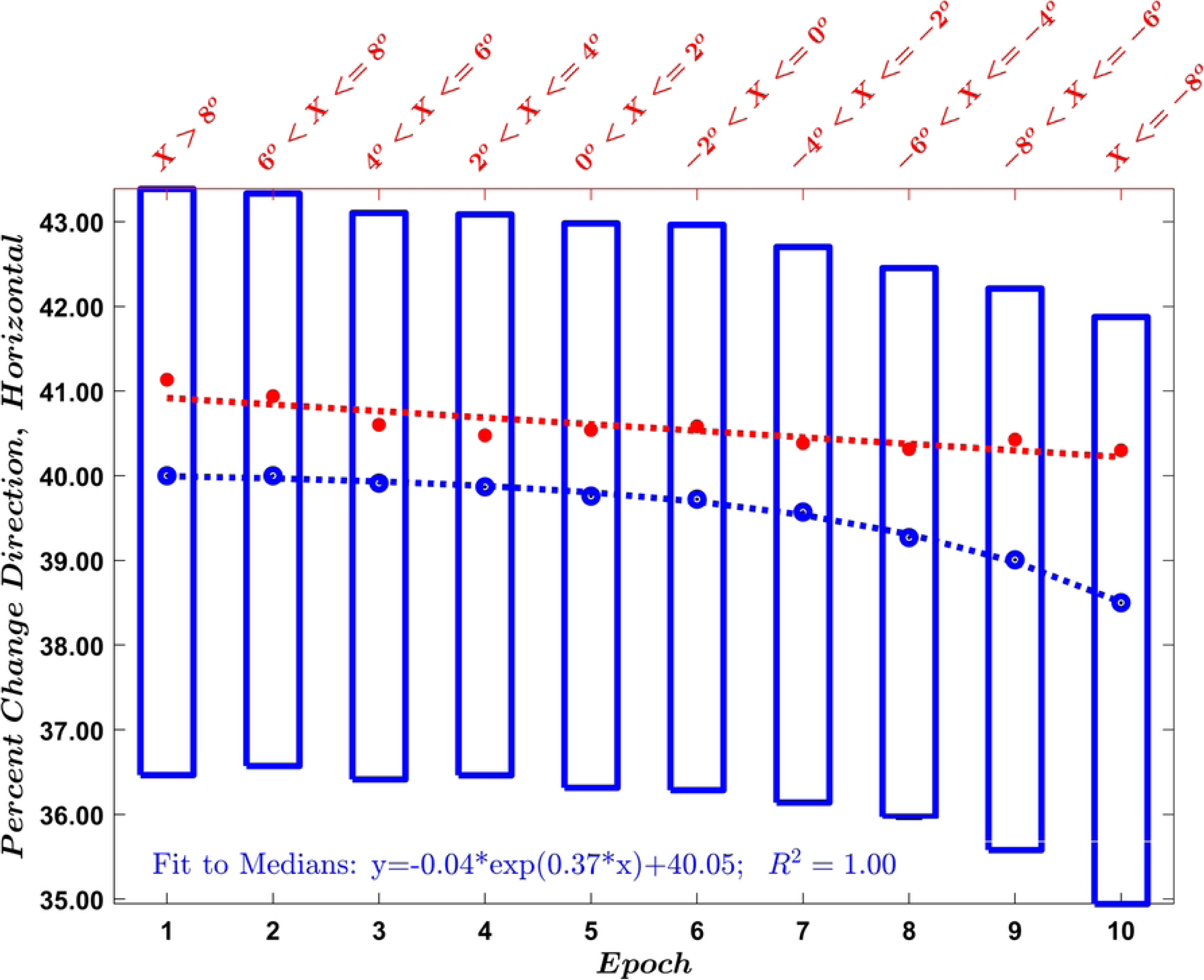
This represents the results of the Kruskal-Wallis test for Mean Intersample Distance (Horizontal), which is a non-parametric analog to the OneWay ANOVA. The *χ̃*^2^ is analogous to the *F* − *value*, with *df* = (9*/*116, 656). The p-value is the probability that these results could have occurred by chance. Post-hoc analyses, controlled for multiple comparisons, indicated that the following epochs were not statistically significant different: [1 vs 2,3,4], [2 vs 3,4], [3 vs 4,5], [4 vs 5,6], [5 vs 6] and [6 vs 7]. All other comparisons were statistically significantly different.

### Percent Change

The results for Percent Change Direction are presented in Fig. 8. The medians of Percent Change Direction (in blue) are relatively stable for epochs one to five. From then on, there is a gradual (exponential) decrease in Percent Change Direction. The fit of the exponential proportional growth rate curve (blue dashed line) is good. The red line evaluates the effect of vertical position during the random saccade task. The trajectory of the red line is declining gradually and more or less linearly from high vertical position to low vertical position. The trajectories for epoch versus Position are nonetheless very different. It seems reasonable to conclude that vertical position may be contributing slightly to the decline across epochs in median Percent Change Direction.

**Fig 8.**
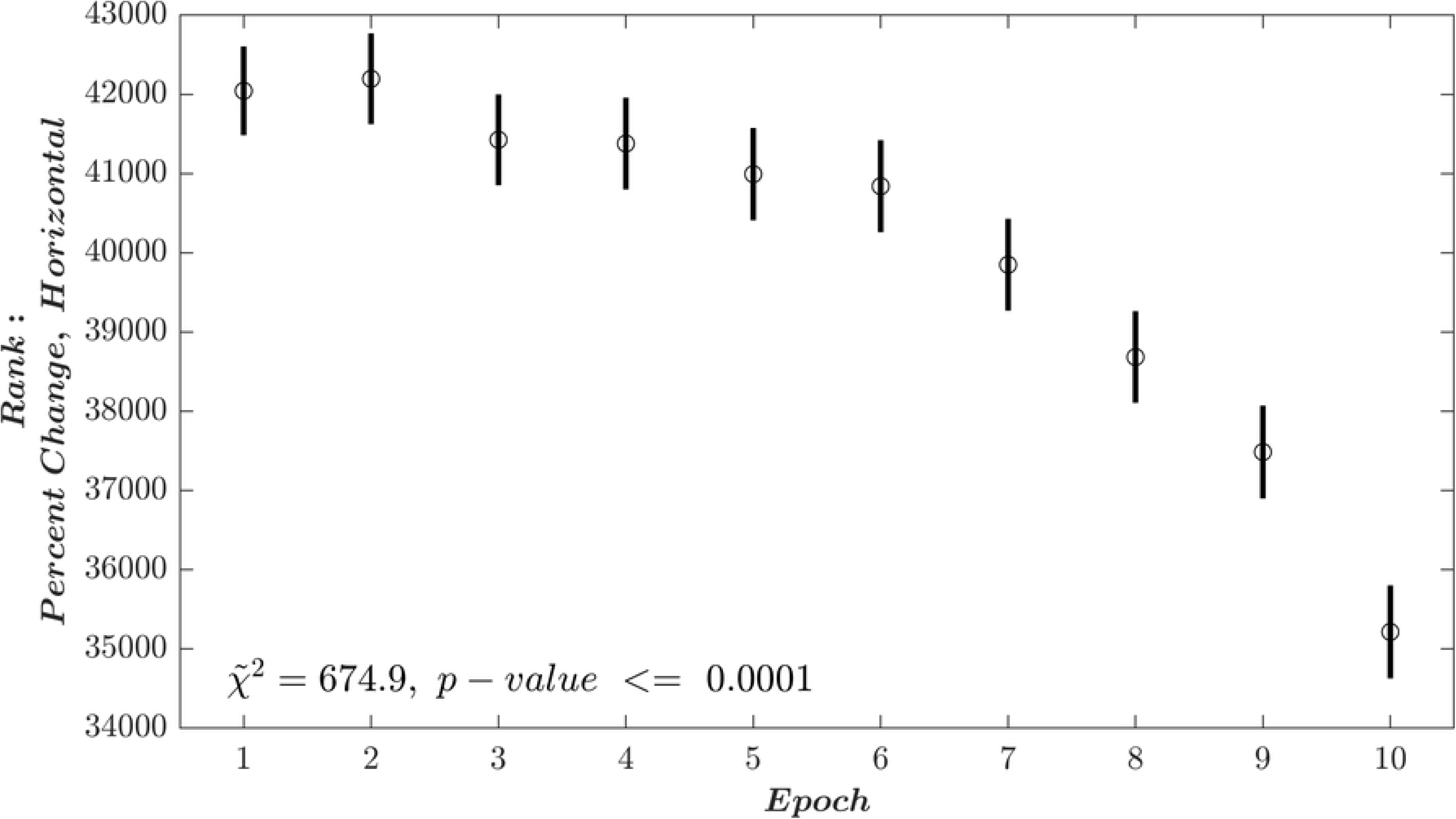
Results for Percent Change (Horizontal). See Fig. 5 for details. The position data (red dots) are fit well to a regression line with a negative slope (*R*^2^ = 0.75, *F* = 24.1, *p − value* = 0.0012, *df* = 1*/*9)

In Fig. 9, we present the statistical results comparing all the Percent Change Directions for all fixations during each epoch. The *χ̃*^2^ is very large, meaning that there are definitely statistically significant differences in Percent Change Direction across epochs.

**Fig 9.**
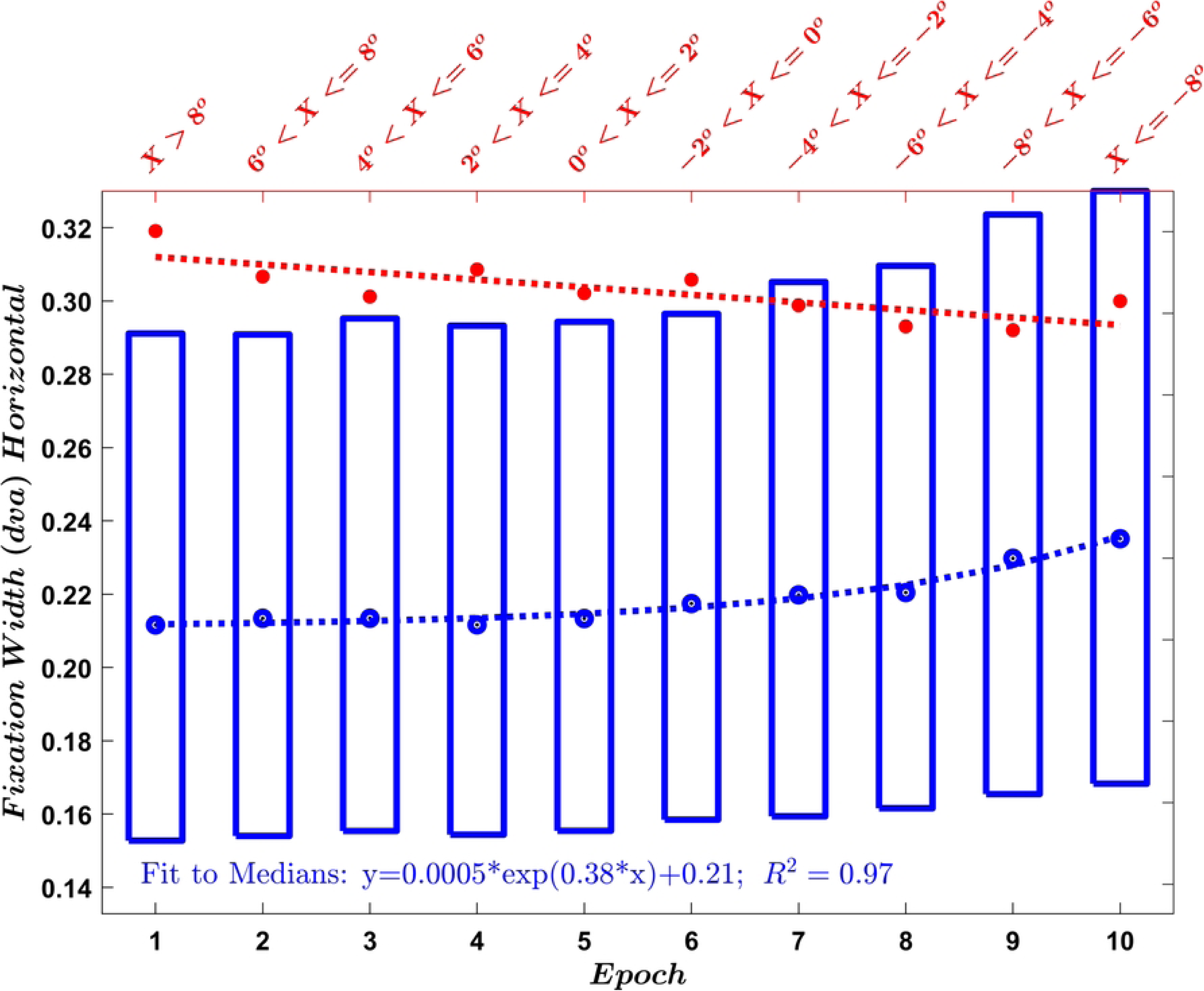
Analysis of Percent Change (Horizontal). See caption for Fig. 6 for details. Post-hoc analyses indicated that the following epochs were not statistically significant different: [1 vs 2,3,4,5], [2 vs 3,4], [3 vs 4,5,6], [4 vs 5,6], [5 vs 6,7] and [6 vs 7]. All other comparisons were statistically significantly different.

### Fixation Width

The results for Fixation Width are presented in Fig. 10. The median Fixation Widths (in blue) are relatively stable for epochs one to five. From then on, there is a gradual (exponential) increase in Fixation Width. The fit of the exponential proportional growth rate curve (blue dashed line) is good. The median number of letters per fixation was 0.54. The letter width was 0.4 dva. So, for epoch 1, the median fixation width was approximately 50% of the letter width. The red line evaluates the effect of vertical position during the random saccade task. The red line is fit well to a regression line with a negative slope. The trajectory of the red line is very different from the trajectory of the blue line. We interpret this to mean the reading task results for Fixation Width are not influenced by vertical position.

**Fig 10.**
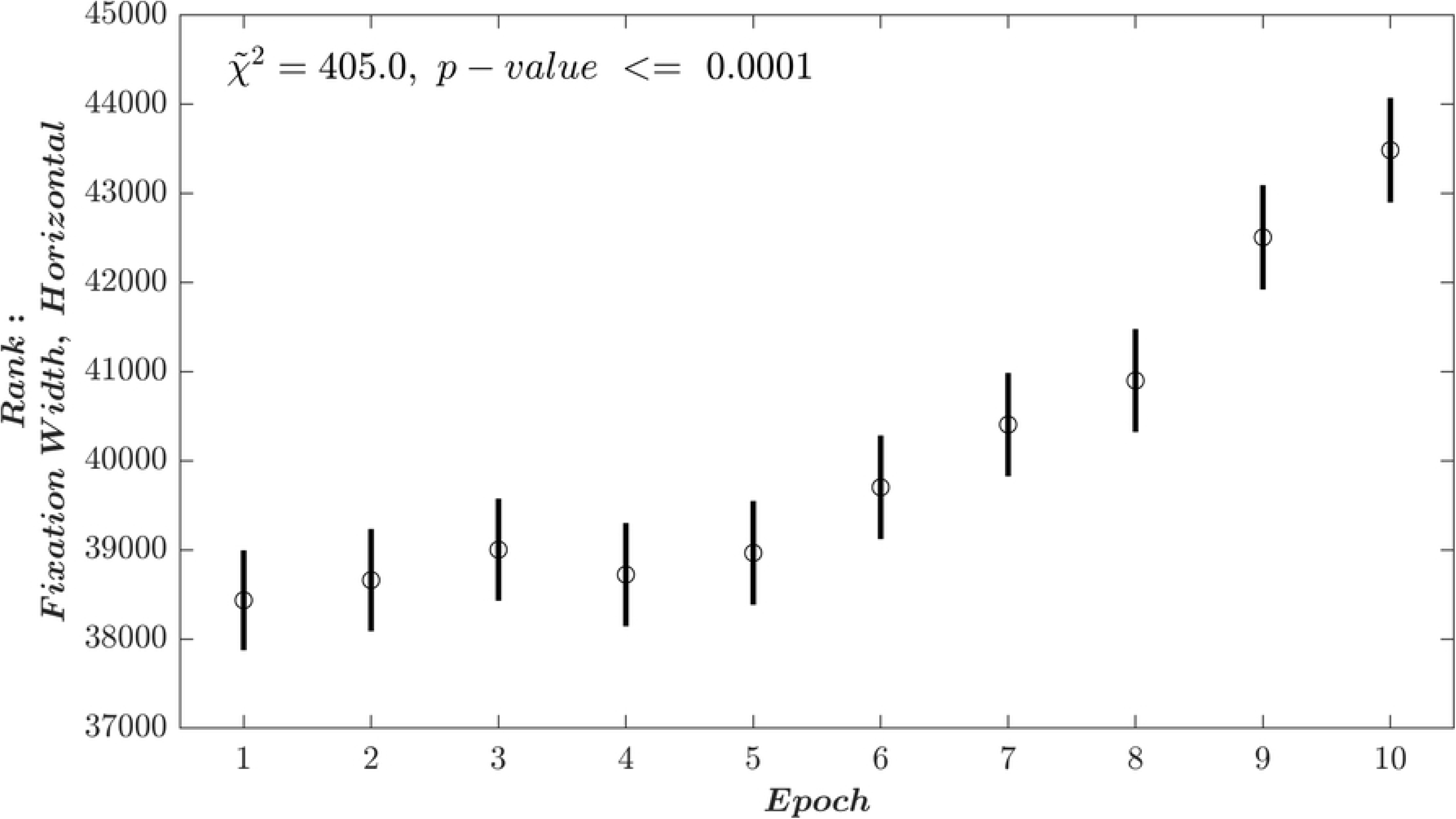
Results for Fixation Width (Horizontal). See Fig. 5 for details. The position data (red dots) are fit well to a regression line with a negative slope (*R*^2^ = 0.62, *F* = 13.5, *p − value* = 0.0063, *df* = 1*/*9)

In Fig. 11, we present the statistical results comparing all the Fixation Widths for all fixations during each epoch. The *χ̃*^2^ is very large, meaning that there are definitely statistically significant differences in Fixation Width across epochs.

**Fig 11.**
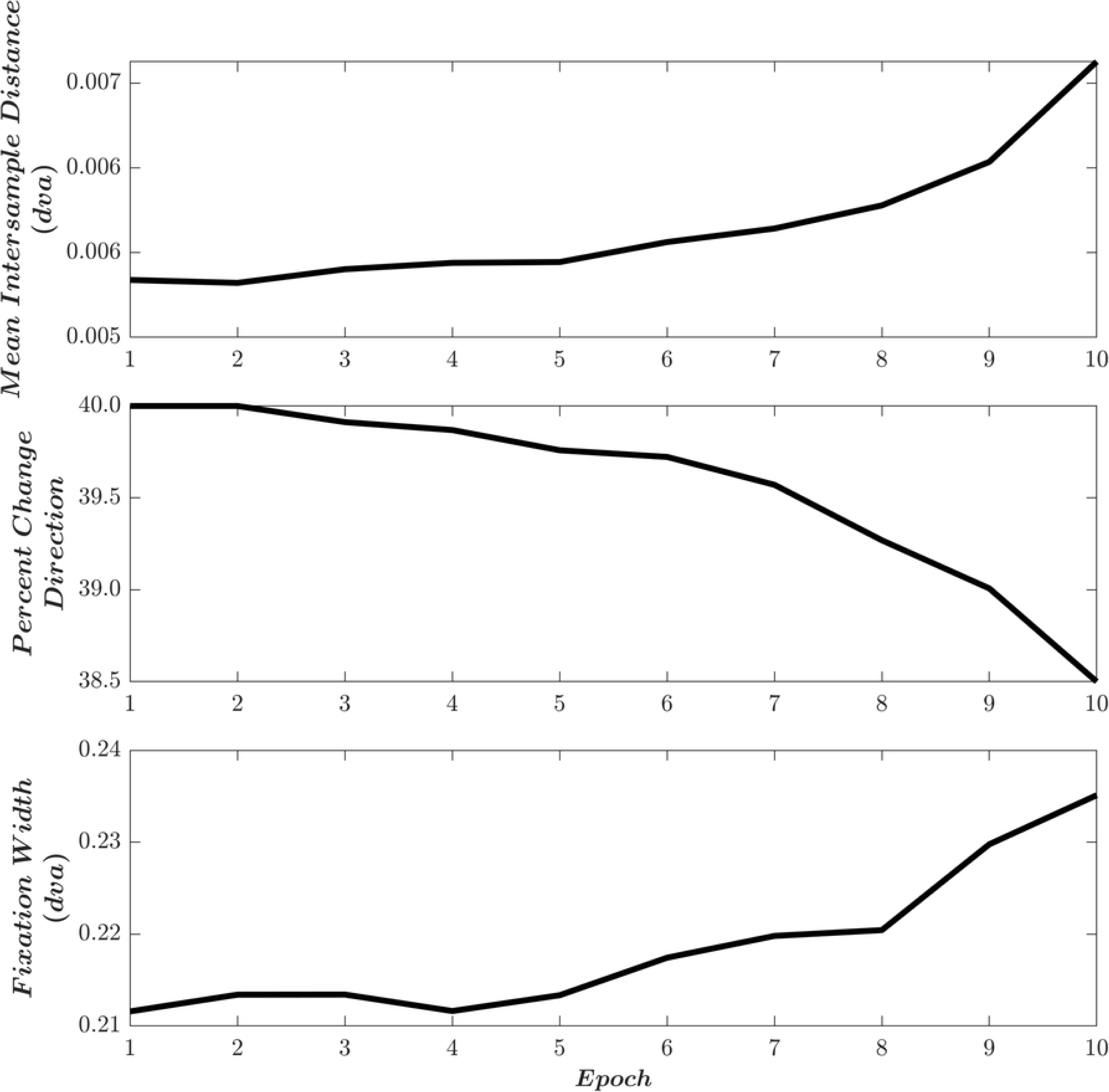
Analysis of Fixation Width (Horizontal). See caption for Fig. 6 for details. Post-hoc analyses indicated that the following epochs were not statistically significant different: [1 vs 2,3,4,5], [2 vs 3,4,5,6], [3 vs 4,5,6], [4 vs 5,6], [5 vs 6], [6 vs 7], [7 vs 8] and [9 vs 10]. All other comparisons were statistically significantly different.

### Inter-Correlations between measures

Table 3 presents the inter-correlations between our three key metrics.

**Table 3.**
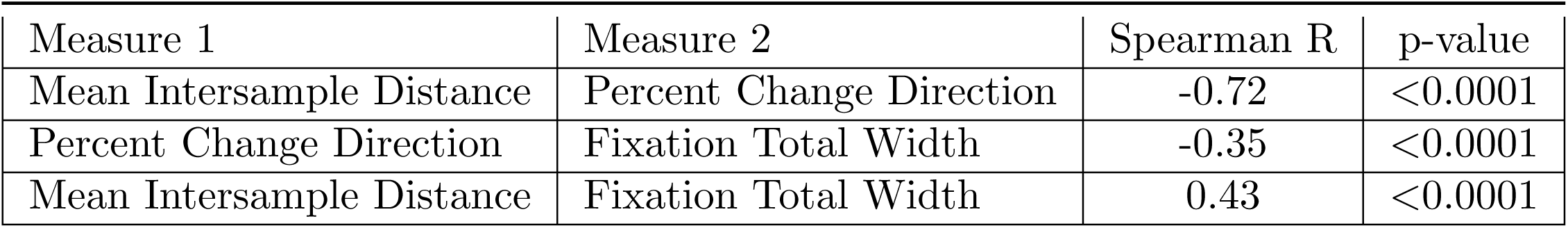
Table of Intercorrelations.

| Measure 1 | Measure 2 | Spearman R | p-value |
| --- | --- | --- | --- |
| Mean Intersample Distance | Percent Change Direction | -0.72 | <0.0001 |
| Percent Change Direction | Fixation Total Width | -0.35 | <0.0001 |
| Mean Intersample Distance | Fixation Total Width | 0.43 | <0.0001 |

### Summary Plot

We present a summary plot of all of the key metrics used in this study. as a function of time-on-task (Fig. 12). The trajectories for Mean Intersample Differences and Fixation Total Width are very similar. As was noted in Table 3, these two metrics are positively correlated. Both Mean Intersample Distance and Fixation Total Width have negative correlations with Percent Change Direction, which has the opposite trajectory of the other two measures.

**Fig 12.**
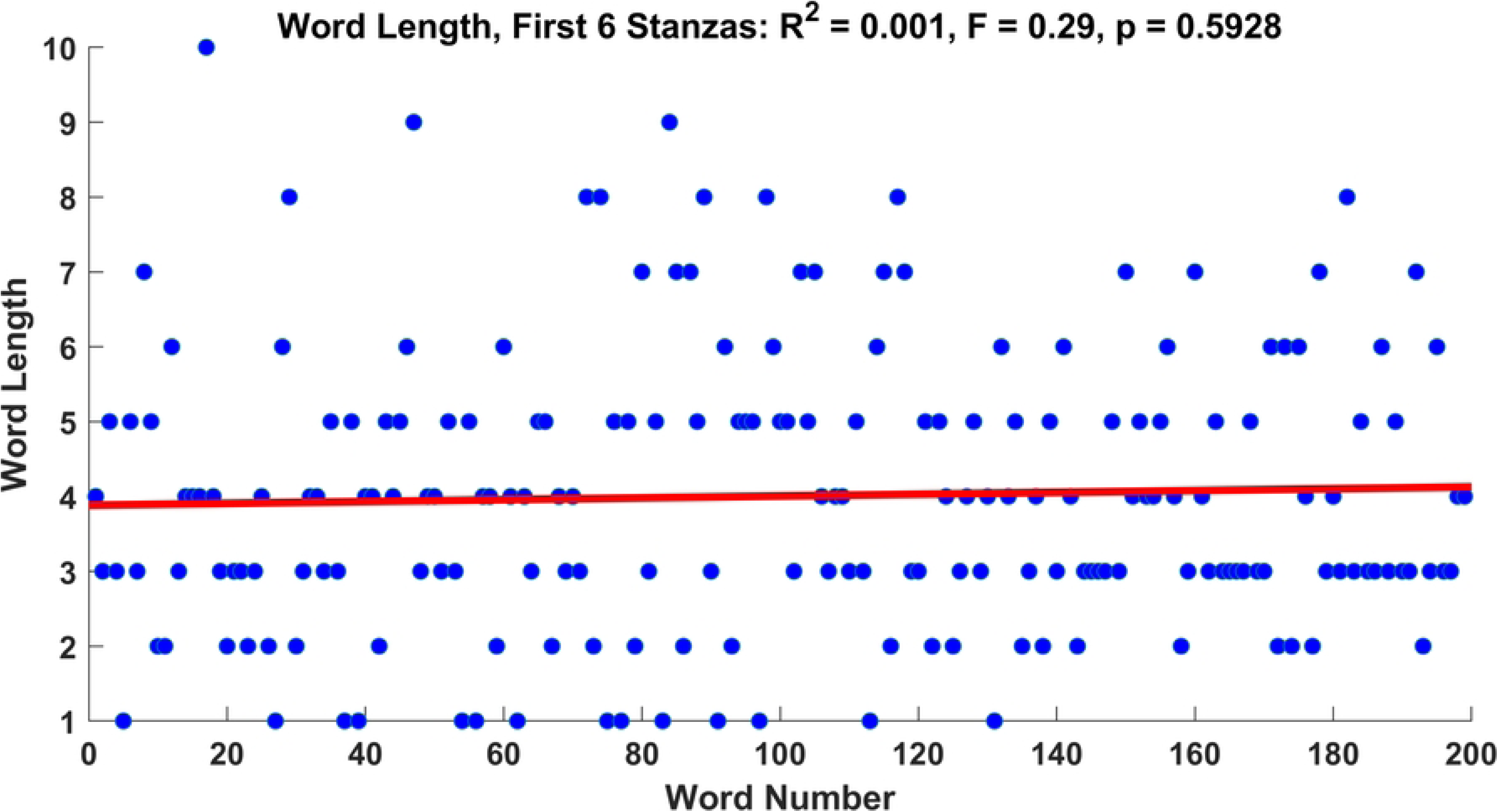
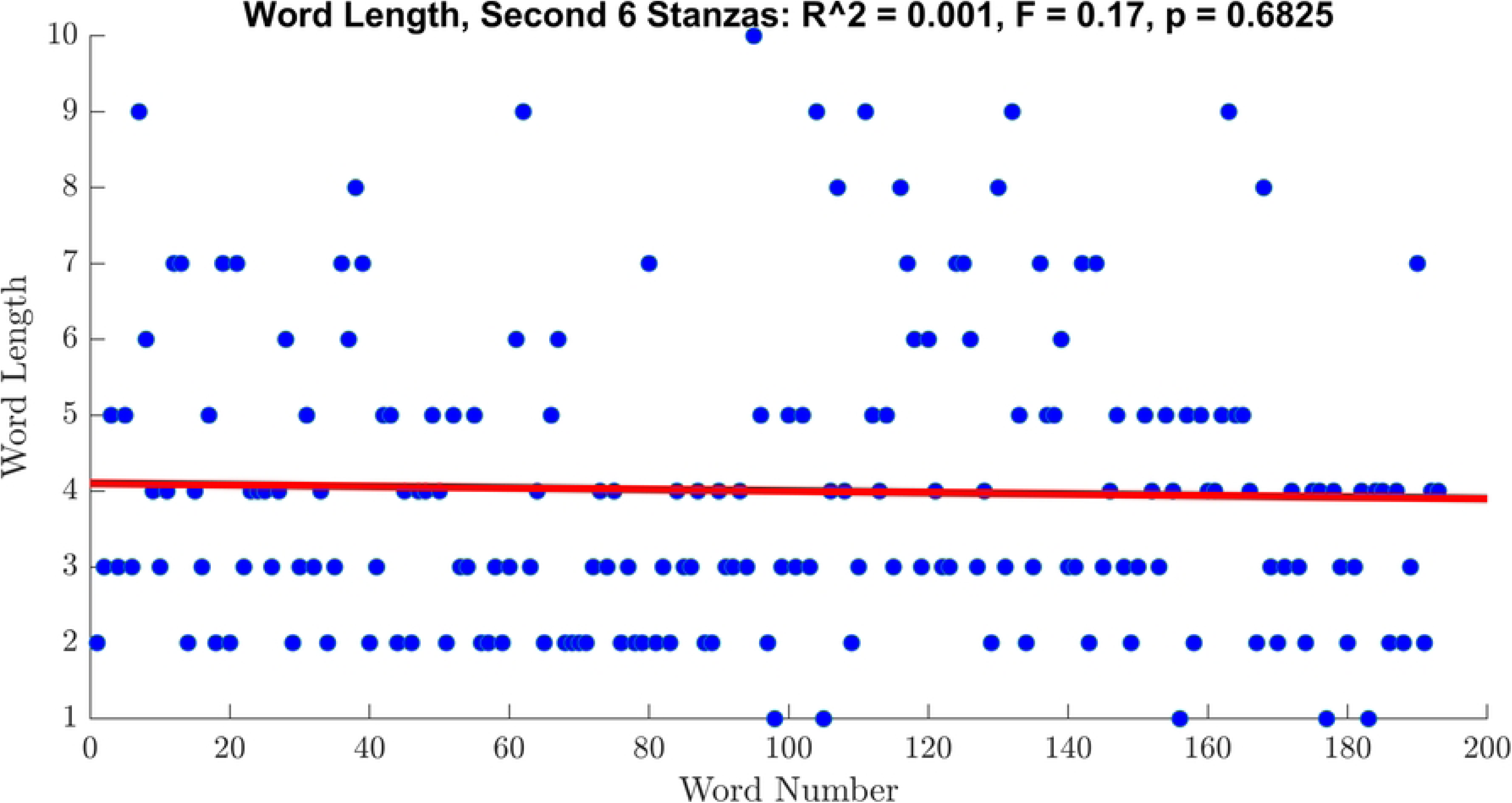
Summary of Results.

### Word Length

Figures 13 and 14 present our analysis of word length. There is no discernible trend across word number for either set of stanzas. Therefore, we conclude that word length is not an important influence on our time-on-task metrics.

**Fig 13.**
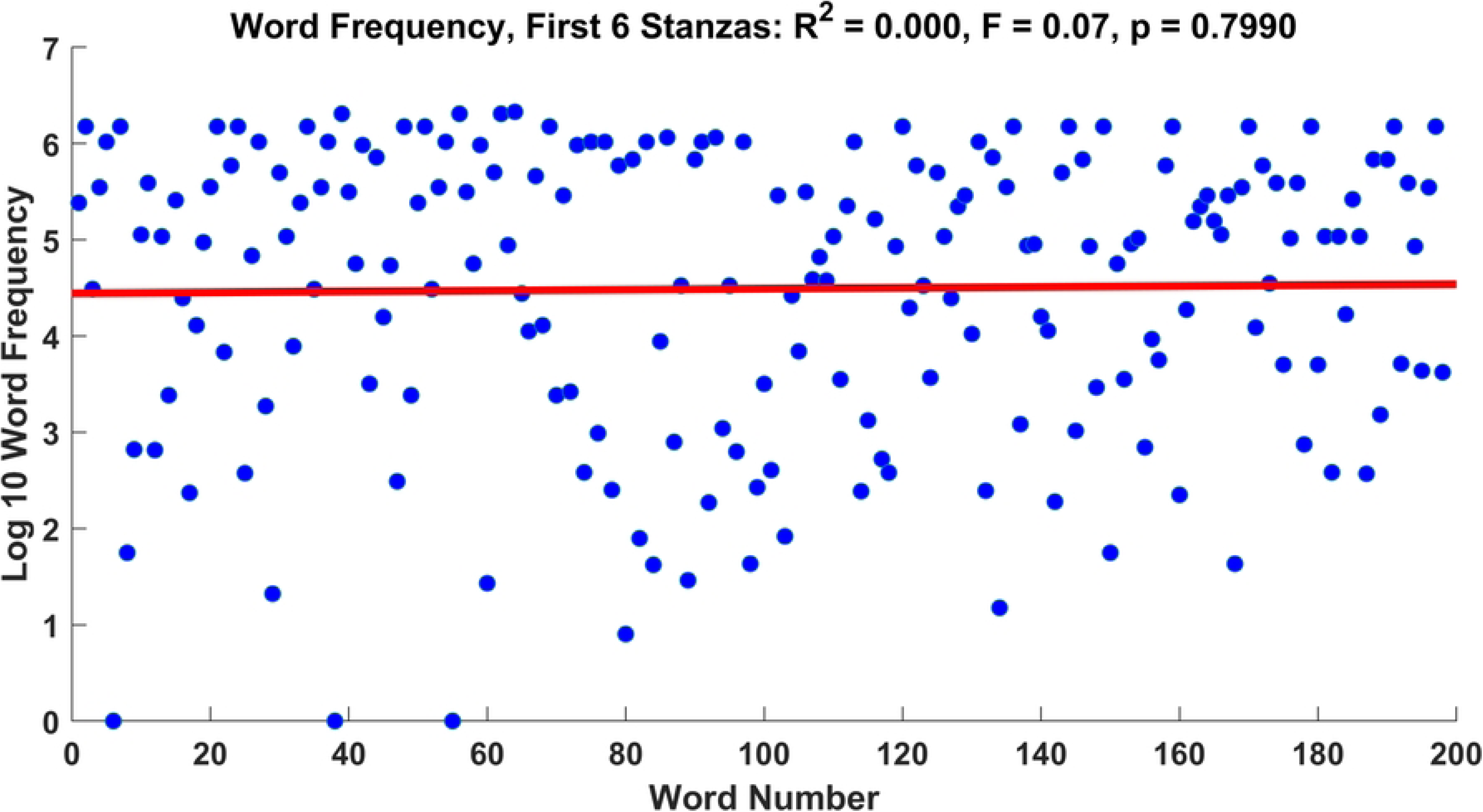
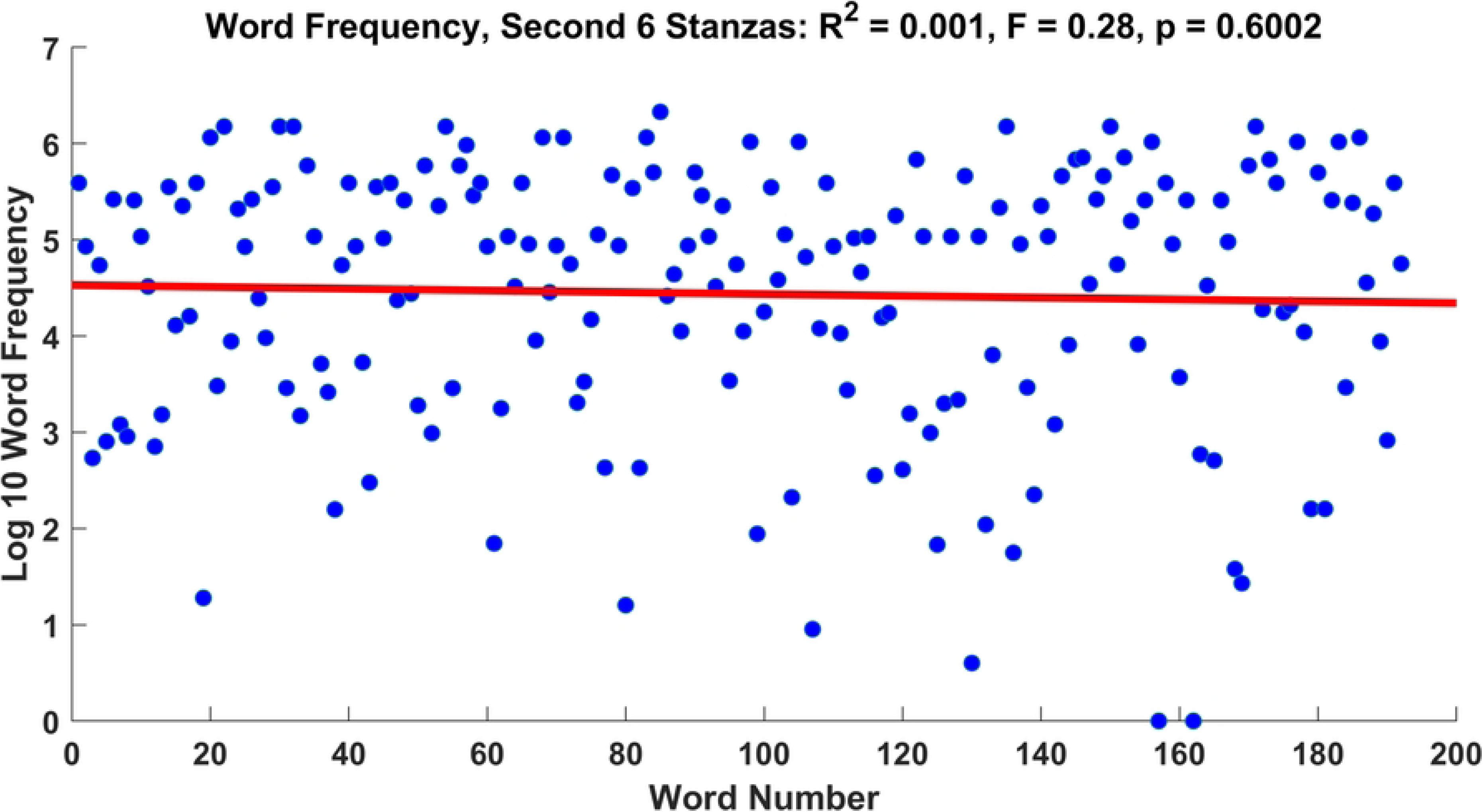
Analysis of word length as a function of word number in the first 6 stanzas of the poem (Appendix Fig. 1).

**Fig 14.**
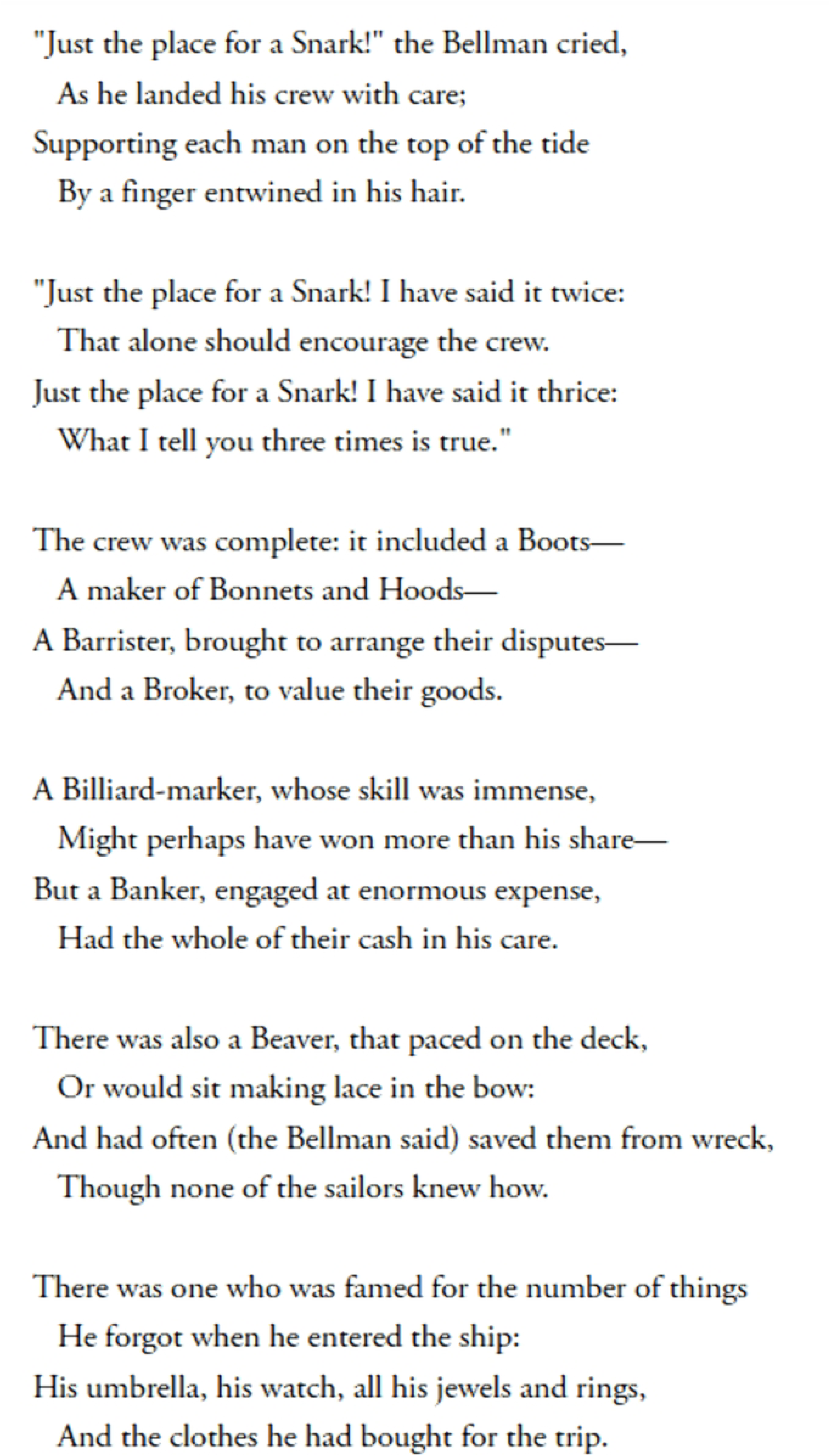
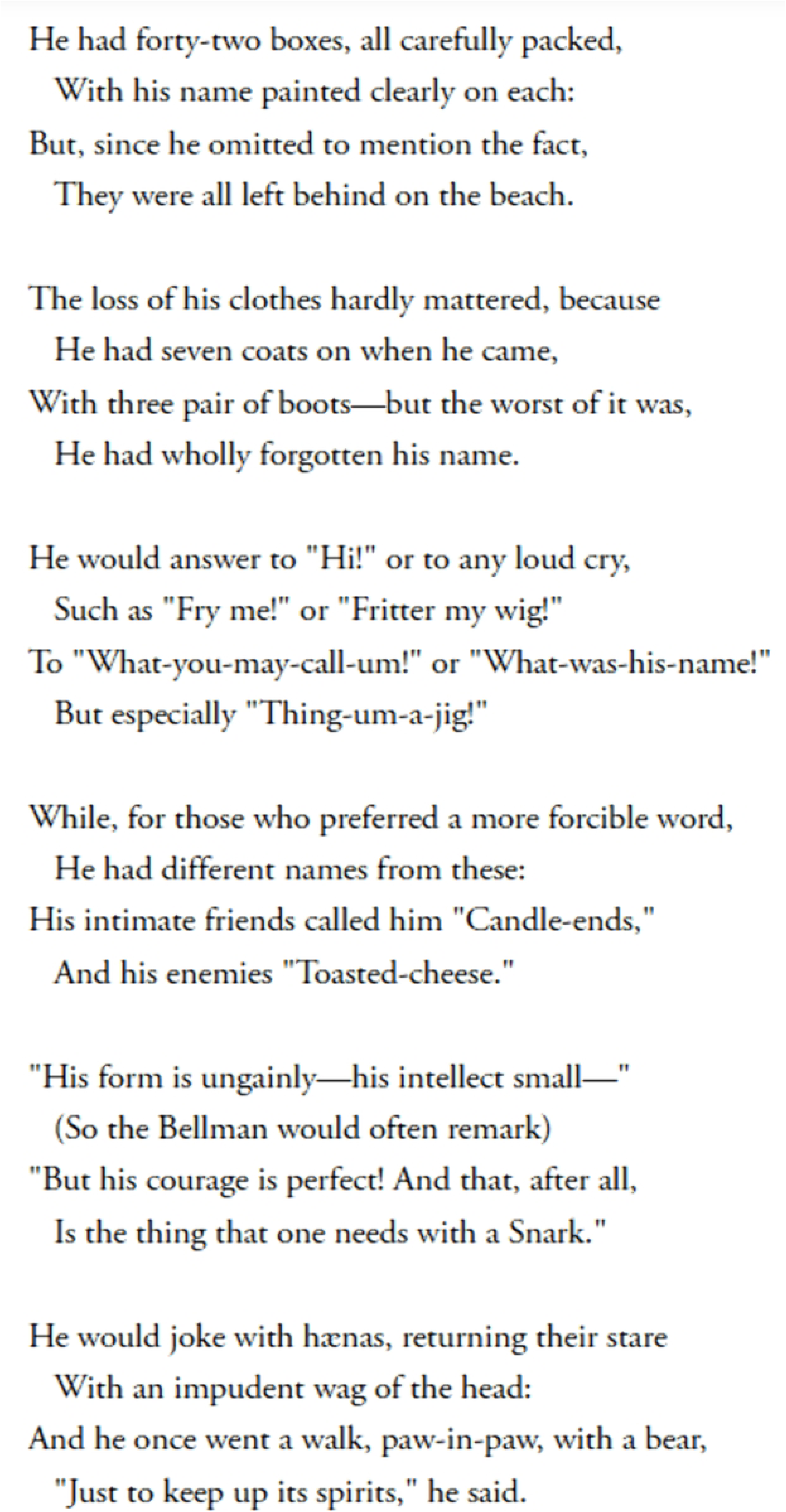
Analysis of word length as a function of word number in the second six stanzas of the poem (Appendix Fig. 2).

### Word Frequency

Figures 15 and 16 present our analysis of word frequency. There is no discernible trend across word frequency for either set of stanzas. Therefore, we conclude that word frequency is not an important influence on our time-on-task metrics.

**Fig 15.**
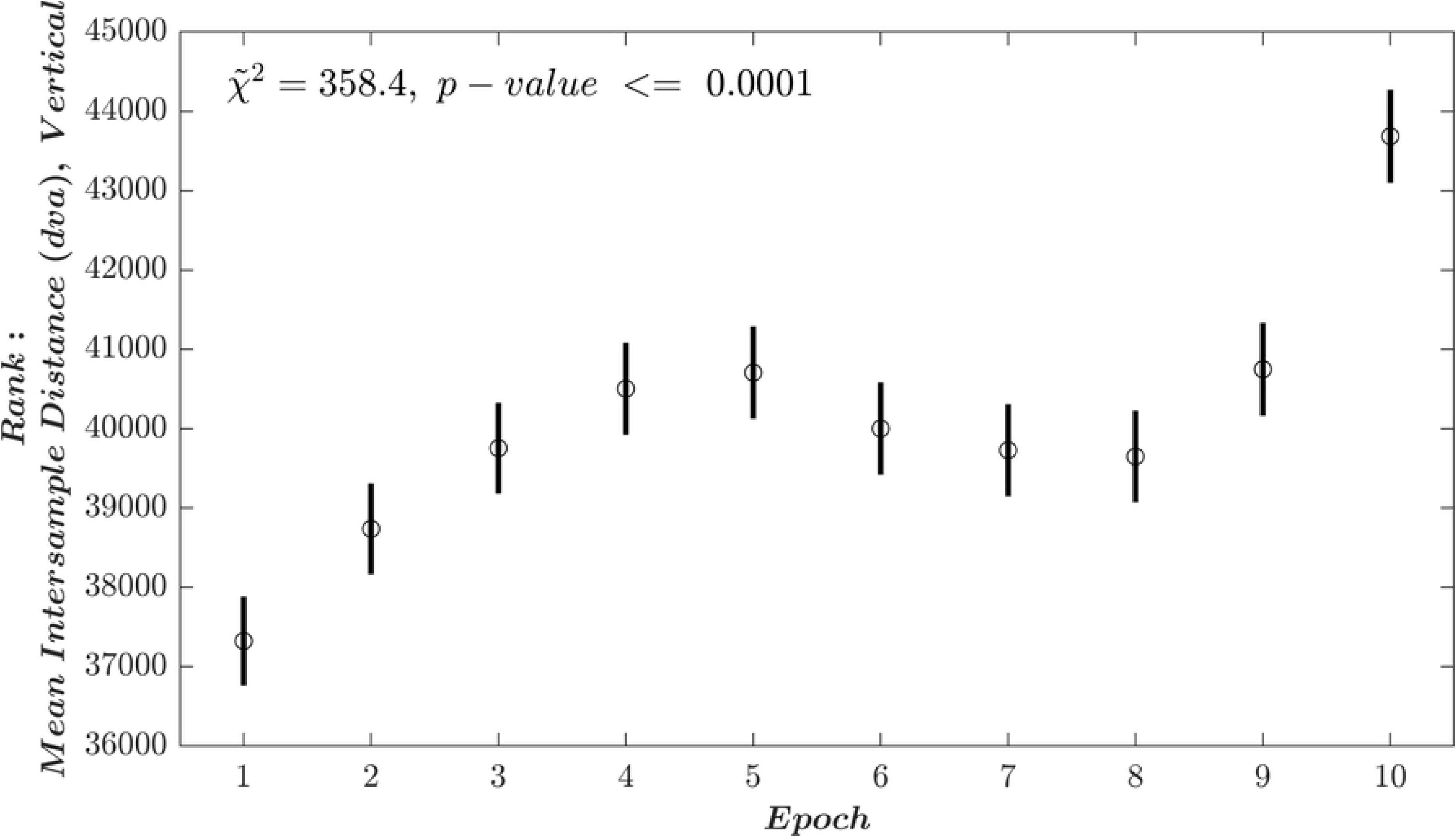
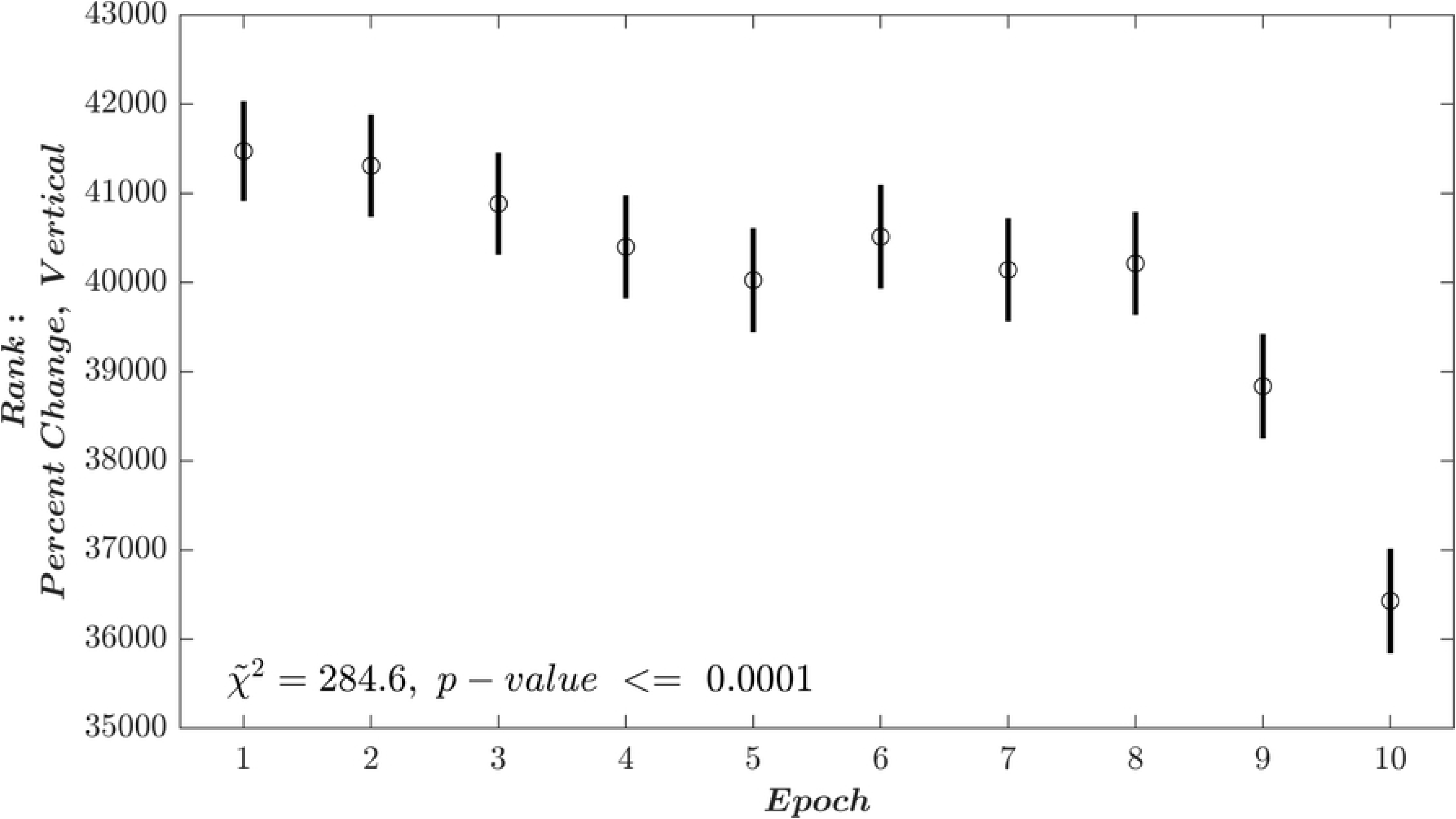
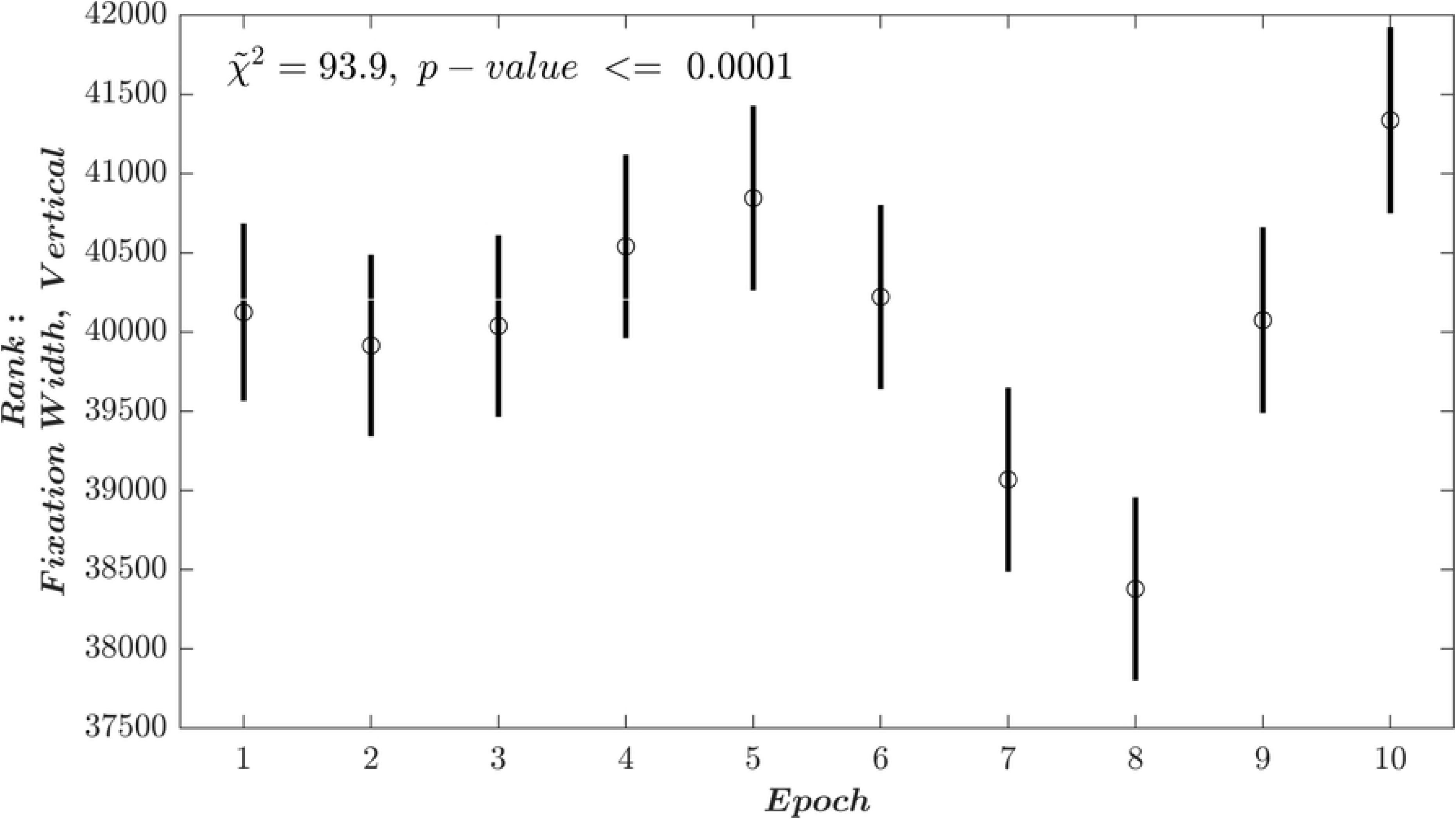
Analysis of word frequency as a function of word number in the first 6 stanzas of the poem (Appendix Fig. 1).

**Fig 16.**
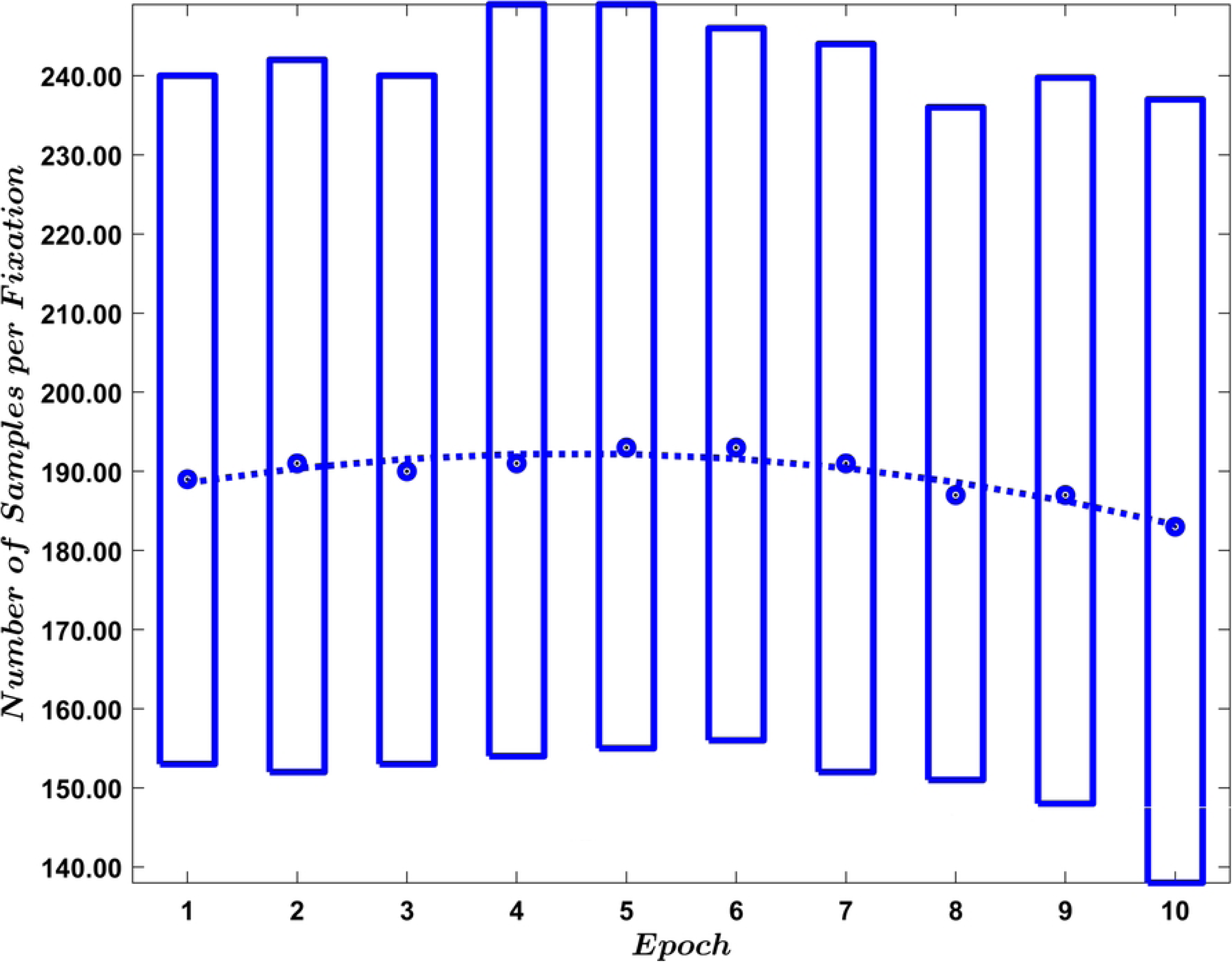
Analysis of word frequency as a function of word number in the second 6 stanzas of the poem (Appendix Fig. 2).

## Discussion

We report changes in ocular fixation characteristics as a function of time on task during poetry reading. The most interpretable results were for horizontal position. The changes across epochs for these measures were generally monotonic. The strongest effect (based on the *χ̃*^2^ value) was the increase in the Mean Intersample Distance (dva). It changed little in the first five epochs but then increased exponentially for epochs six to ten. Another strong effect was Percent Change Direction which decreased exponentially over the 10 epochs. These two measures were highly inter-correlated (Table 3 (−0.72)). Fixation Width also increased exponentially across epochs. These effects were unrelated to changes in word length or word frequency. We are not aware of another study that looked at changes in fixation characteristics during reading as a function of time on task.

The best summary of our results are illustrated in Figs. 1 and 2. The first figure is an archetypal (and perhaps extreme) illustration of fixation during the start of the reading task. The fixation is narrow in the horizontal dimension, with small Mean Intersample Differences and frequent direction changes. The second figure is an archetypal illustration of fixation during the end of the reading task. The fixation is wide overall, the Mean Intersample Differences are large and there are many fewer direction changes. Another good summary plot is Fig. 9, which illustrates the changes across metrics as reading progresses. In all cases, the metrics are relatively stable until the sixth epoch at which point exponential changes appear.

Perhaps one way to think about these changes with time on task is that, as reading progresses, it becomes more difficult to hold the eye still. Early in reading, the total width is lower, the mean intersample distance is lower and there are frequent adjustments (i.e., changes in direction) to keep the eye well focused on the target word. As time progresses, it becomes more and more difficult to hold the eye perfectly steady, so fixations become wider and mean intersample distance increases.

As noted above, with our task, time-on-task is confounded with vertical position. Theoretically, the changes we observed could be affected by vertical position rather than time-on-task. For all metrics, there were statistically significant linear changes as a function of vertical position in the random saccade task. In two cases (Mean Intersample Difference and Percent Change Direction), the change as a function of vertical position was in the same direction as the change across epochs (time). However, the trajectories were distinct from our epoch data. We think it is reasonable to conclude that our time-on-task (epoch) data were not simply a reflection of vertical position. In the case of Fixation Width, the trajectories move in opposite directions and therefore we assume that our fixation width results are not simply a reflection of vertical position.

Future studies might be designed to minimize or eliminate the issue of vertical position. Text could be presented in the middle of the screen, and subjects could indicate in some manner when to display another stanza, for example. If the text were displayed in the middle of the screen we would not need to assess the confounded effects of changes in vertical position.

Although we found statistically significant changes in our metrics for vertical position, the results were not monotonic. We have no explanation of these complex relationships and so we de-emphasize them in this report. Future studies could be designed to explore these relationships.

## Conclusion

## Appendix

Fig 1. *(Appendix)* First six stanzas of the poem presented during session 1.

Fig 2. *(Appendix)* Second six stanzas of the poem presented during session 2.

Fig 3. *(Appendix)* Analysis of Mean Intersample Distance (Vertical). See caption for Fig. 4 in the main document for details. Post-hoc analyses indicated that the following epochs were not statistically significant different: 2 vs 3,7,8], [3 vs 4,5,6,7,8,9], [4 vs 5,6,7,8,9], [5 vs 6,7,8,9], [6 vs 7,8,9], [7 vs 8,9] and [8 vs 9]. All other comparisons were statistically significant.

Fig 4. *(Appendix)* Analysis of Percent Change (Vertical). See caption for Fig. 4 in the main document for details. Post-hoc analyses indicated that the following epochs were not statistically significant different: [1 vs 2,3,4,6], [2 vs 3,4,6,8], [3 vs 4,5,6,7,8], [4 vs 5,6,7,8], [5 vs 6,7,8], [6 vs 7,8] and [7 vs 8]. All other comparisons were statistically significantly different.

Fig 5. *(Appendix)* Analysis of Fixation Width (Vertical). See caption for Fig. 4 in the main document for details. Post-hoc analyses indicated that the following epochs were not statistically significant different: [1 vs 2,3,4,5,6,7,9], [2 vs 3,4,5,6,7,9], [3 vs 4,5,6,7,9], [4 vs 5,6,9,10], [5 vs 6,9,10], [6 vs 7,9,10] and [7 vs 8,9].

